# JUNO-Coated Beads as a Functional Assay to Capture and Characterize Fertilization-Competent Human Sperm

**DOI:** 10.1101/2025.09.30.679507

**Authors:** Paula Cots-Rodríguez, Mirian Sanchez-Tudela, Julieta G. Hamze, Patrick Yip, Emilio Gómez, Jeffrey E. Lee, María Jiménez-Movilla

**Affiliations:** Department of Cell Biology and Histology, Faculty of Medicine, CEIR Campus Mare Nostrum (CMN), University of Murcia, Murcian Institute of Biosanitary Research Pascual Parrilla (IMIB), Health Sciences Campus, 30120 Murcia, Spain; Department of Laboratory Medicine and Pathobiology, Temerty Faculty of Medicine, University of Toronto, M5S 1A8 Toronto, ON, Canada; IVF Laboratory, Next Fertility /Gametia, Murcia, Spain

**Keywords:** Fertilization, spermatozoa selection, spermatozoa vitrification, in vitro sperm binding model, human-based model, fertilization-competent spermatozoa, acrosome reaction, DNA integrity, JUNO, IZUMO1

## Abstract

**Study question:** Can human fertilization-competent spermatozoa be captured through their ability to bind the oocyte receptor JUNO?

**Summary answer:** JUNO-coated beads, which mimic the oocyte geometry, selectively bound acrosome-reacted spermatozoa with intact DNA, revealing that vitrification preserves functional sperm binding while slow cryopreservation increases non-specific interactions.

**What is known already:** It is well established that sperm must undergo the acrosome reaction and expose the receptor IZUMO1 on the sperm head to bind specifically to JUNO on the oolemma. Studying the spermatozoa that reaches and engages with the oolemma, however, remains highly challenging due to the technical difficulty of recovering these sperm at the site of molecular interaction. Bead-based models that content oocyte receptors have therefore emerged as a powerful approach to functionally assess sperm-oocyte interactions, with promising applications for evaluating sperm quality.

**Study design, size, duration:** This was a cross-sectional experimental study including 21 semen donors of reproductive age recruited between January 2023 and June 2025. The JUNO-bead-based model was first validated using fresh human semen samples to establish optimal sperm concentration and co-incubation time. Subsequently, two semen preservation methods, slow freezing and rapid freezing, were compared with respect to sperm binding capacity to JUNO-coated beads, acrosomal status, and DNA integrity. Finally, donors were classified according to sperm binding capacity.

**Participants/materials, setting, methods:** Recombinant JUNO protein was expressed and purified in Drosophila melanogaster S2 cells, and protein-bead conjugation was verified by immunochemistry. Human semen samples were obtained from donors aged 19-42 years, including both fresh ejaculates and cryopreserved samples. Sperm binding capacity, acrosome reaction, and DNA fragmentation were assessed using widefield fluorescence microscopy and flow cytometry. Specificity of sperm-bead binding was evaluated with anti-IZUMO1 monoclonal antibodies.

**Main results and the role of chance:** Human JUNO recombinant protein was successfully conjugated to oocyte-sized beads to generate a sperm-binding assay mimicking the geometry of the oocyte and experimental conditions of the *in vitro* fertilization. Human sperm bound specifically to JUNO-beads in a dose- and time-dependent manner, with highly significant differences compared to control beads (*p* ≤ 0.0001). Vitrified-based cryopreserved sperm displayed higher binding to JUNO-beads than conventionally cryopreserved samples (*p* ≤ 0.0001). Binding was significantly inhibited by an anti-IZUMO1 (2.5 ug/mL) antibody that blocks specifically the IZUMO1-JUNO interaction in vitrified samples (p ≤ 0.01), but not in conventionally cryopreserved sperm. Sperm bound to JUNO-beads were predominantly acrosome-reacted in both preservation methods; however, vitrified samples retained higher DNA integrity compared with conventionally cryopreserved samples. The assay proved robust across multiple donors and ejaculates, allowing classification into low- and high-binding capacity (LBC and HBC) groups. Pearson correlation analyses revealed only weak associations between total sperm motility and bead-binding parameters (|r| < 0.27), indicating negligible or absent linear relationships.

**Large-scale data:** N/A

**Limitations, reasons for caution:** This study was performed *in vitro*, and the number of semen donors was limited. As all participants were healthy donors, the population represents a selected fertile subpopulation. Further studies using samples from diverse patient populations are required to validate the potential of the assay as a predictor of male fertility.

**Wider implications of the findings:** This study positions the JUNO-bead binding assay as a powerful functional model to investigate the biology of fertilization-competent sperm. By selectively capturing spermatozoa that have undergone the acrosome reaction and maintain DNA integrity, the model provides a unique experimental platform to study the molecular determinants of fertilization, to refine the selection of sperm for assisted reproduction, and to identify potential targets for novel contraceptive strategies. Beyond preservation protocols, these findings provide new functional evidence that sperm preservation method directly influences the molecular integrity required for fertilization, supporting vitrification as a superior approach over slow freezing. Moreover, the JUNO-bead assay emerges as a sensitive tool to reveal differences in sperm quality that are not captured by standard semen analysis, with potential applications in the optimization of assisted reproduction and fundamental research on the mechanisms that define the fertilizing spermatozoon.

**Study funding/competing interest(s):** This work is part of the project PID 2020-114109GB-I00 funded by MCIN/AEI/10.13039/501100011033 to M.J.M. This work was also supported, in part, by the Gates Foundation [INV-055841]. The conclusions and opinions expressed in this work are those of the author(s) alone and shall not be attributed to the Foundation. Under the grant conditions of the Foundation, a Creative Commons Attribution 4.0 License has already been assigned to the Author Accepted Manuscript version that might arise from this submission. Protein production and biophysics infrastructure is supported by funding from Canada Foundation for Innovation John R Evans Leaders Fund (CFI-JELF) to J.E.L. The authors declare no conflict of interest.

**Lay summary:** Identifying and studying the sperm cell that is truly capable of fertilizing an egg is a major challenge, because this process occurs precisely when the sperm binds to the egg and penetrates it. In this study, we developed a model based on microscopic beads that mimic the shape of the egg and are coated with JUNO, a receptor essential for sperm–egg recognition. These JUNO-beads capture only those sperm cells with the right molecular and cellular properties to bind to the egg membrane and enable fertilization.

Using this system, we compared sperm samples preserved through slow freezing (cryopreservation) or rapid freezing (vitrification). We observed that vitrified samples retained a higher proportion of sperm with fertilizing characteristics, sperm that had undergone the acrosome reaction but maintained intact DNA. Furthermore, by applying this model to samples from different donors, we were able to classify them according to their high or low binding capacity to JUNO-beads.

Overall, this approach provides a new way to “capture” and evaluate fertilizing sperm, offering potential applications for improving sperm quality assessment in assisted reproduction, and a valuable tool for studying the defining features of the fertilizing sperm cell.

## Introduction

The fertilization process relies on a well-coordinated recognition mechanism between two highly specialized haploid cells: spermatozoon and oocyte (Okabe, 2013). Over the past years, several molecular factors involved in gamete interaction and fusion have been identified. On the sperm side, IZUMO1 plays a key role in the fusion process, as demonstrated by IZUMO1 knockout model, where Izumo1^-/-^ spermatozoa, despite their ability to penetrate the zona pellucida fail to bind and fuse with the oolemma (Inoue *et al*., 2005; Matsumura *et al*., 2021). A similar phenotype has been described for DCST1, DCST2, FIMP, SOF1, SPACA6, TMEM81 and TMEM95 proteins, where each knockout model exhibited spermatozoa accumulating in the perivitelline space with impaired fusion ability with oolemma (Lorenzetti *et al*., 2014; Barbaux *et al*., 2020; Fujihara *et al*., 2020; Lamas-Toranzo *et al*., 2020; Noda *et al*., 2020; Inoue *et al*., 2021; Deneke *et al*., 2024).

Conversely, only two factors have been identified on the oocyte side; CD9 and JUNO. Notably, the interaction between sperm IZUMO1 and egg JUNO represents the only well-documented interaction between human gametes (Bianchi *et al*., 2014; Aydin *et al*., 2016a; Kato *et al*., 2016; Ohto *et al*., 2016; Jean *et al*., 2019). The crucial role of JUNO in fertilization was demonstrated using *Juno*^−/−^ oocytes, which allowed penetration of the zona pellucida by wild-type sperm but failed to support binding or fusion with the oolemma (Bianchi *et al*., 2014). Similarly, in experiments using human zona-free oocytes, the presence of an anti-hJUNO antibody prevents human sperm from fusing with and fertilizing the egg (Jean *et al*., 2019).

In this context, from ejaculation to fertilization, spermatozoa undergo physiological transformations, including capacitation, acrosome reaction, and crucial relocation of IZUMO1 protein, all essential for successful interaction with JUNO and thus, fertilization (Yanagimachi, 1994; Inoue *et al*., 2005; Puga Molina *et al*., 2018). Nevertheless, the identification and analysis of fertilization-competent spermatozoa remain technically challenging. Current human sperm characterization methods primarily assess viability, morphology and motility (World Health Organization, 2021). However, a significant proportion of male infertility cases remain idiopathic or unexplained (Ventimiglia *et al*., 2021; Corsini *et al*., 2023), underscoring the need for novel techniques to evaluate the fertilization capacity of apparently normal spermatozoa based on their molecular ability to bind the oocyte.

As a complementary approach to assisted reproductive technologies (ART), sperm cryopreservation represents a crucial strategy for preserving male fertility. To ensure the effectiveness of this technique, it is essential to maintain both sperm viability and functionality after thawing (World Health Organization, 2021). Currently, two main sperm freezing methods exist: slow freezing and rapid freezing. Hereafter, slow freezing will be referred to as *cryopreservation*, and rapid freezing as *vitrification*. These methods differ in cooling rates, cryoprotectant composition, sample volume, storage devices, and thawing protocols (Li *et al*., 2019; Tao *et al*., 2020). Although vitrification has shown promising results, further evaluation of its impact on sperm fertilization capacity is required.

One of the major challenges in assessing the fertilization potential of both fresh and preserved sperm is the limited availability of human oocytes for *in vitro* fertilization (IVF) assays. For this reason, alternative methodologies have been developed to evaluate the sperm-oocyte binding capacity. The human zona pellucida binding assay tests sperm fertilizing potential by measuring their ability to bind to isolated human zonae pellucidae, showing a correlation with *in vitro* fertilization (IVF) outcomes (Burkman *et al*., 1988; Liu, 2003). The zona-free hamster oocyte penetration test evaluates sperm function by testing their ability to undergo capacitation, acrosome reaction, and fuse with the oolemma (Yanagimachi *et al*., 1976). However, due to its lack of physiological relevance, it may yield false negatives, as some sperm that fail this assay can still fertilize human oocytes *in vitro* and *in vivo* (“Group Discussion,” 1986).

Given these limitations, both tests have been classified as obsolete in the latest edition of World Health Organization (WHO) Laboratory Manual for the Examination of Human Semen (World Health Organization, 2021). To address these shortcomings, a three-dimensional (3D) *in vitro* model of sperm-oocyte interaction has been developed (Hamze *et al*., 2020a, Hamze *et al*., 2020c, Hamze *et al*., 2020b). This system uses magnetic sepharose microbeads (hereafter referred to as *beads*) coated with recombinant oocyte proteins, such as zona pellucida (ZP) proteins (Hamze *et al*., 2020b) and JUNO (Hamze *et al*., 2020c), to mimic the size, shape, and molecular surface characteristics of the oocyte. The model has been previously validated in non-human species, including bovine and porcine. In the present study, we optimized a bead-based sperm binding assay specifically for human sperm by employing microbeads coated with recombinant human JUNO. This approach allows for the evaluation of sperm binding capacity without the need for live human oocytes. Once validated, the model was applied to compare the fertilizing potential of spermatozoa preserved using cryopreservation versus vitrification techniques.

## Material and Methods

### 1. Ethical Approval

This study was approved and supervised by the Research Ethical committee (CEI 3094/2020) and Experimental Biosecurity committee (CBE 371/2020 and CBE 576/2023) of Universidad de Murcia.

### 2. Recombinant Protein Design and Expression

The expression and purification of recombinant human JUNO ectodomain was performed following the protocol previously described in (Aydin *et al*., 2016b). The gene corresponding to full-length human JUNO (GenBank accession number: NM_001199206, residues 1-250) was codon optimized for expression in *Drosophila melanogaster* and gene synthesized (Integrated DNA Technologies, USA). The DNA sequences encoding the extracellular regions of JUNO (residues 20-228) with a BiP signal peptide were subcloned into a modified pMT-puromycin expression vector (Invitrogen, USA). The protein constructs contain a C-terminal thrombin cleavage site and 10X-His affinity tag, except in the control group, where an untagged construct was used. The JUNO20-228 pMT expression plasmid was stably transfected in *Drosophila* S2 cells (Invitrogen, USA) using Effectene transfection reagent (Qiagen, Germany), according to the manufacturer’s protocol. Briefly, Drosophila S2 cells were cultured in Schneider’s medium (Gibco, Thermo Fisher Scientific, USA) supplemented with 10% (v/v) heat-inactivated fetal bovine serum (FBS) plus 1X antibiotic-antimycotic (Gibco, Thermo Fisher Scientific, USA) and propagated at 27 °C. 2 μg expression plasmid was mixed with the transfection reagents and the transfection complexes were added drop-wise onto the S2 cells. At 72 hours post-transfection, the cultured media were replaced with fresh S2 growth media supplemented with 6 μg/mL puromycin (Bioshop, Canada). Subsequently, S2 cells were gradually adapted to serum-free Insect-XPRESS growth media (Lonza, Switzerland) with 6 μg/mL puromycin. Stably transfected cells were grown to 1×10^7^ cells/mL in Insect-XPRESS growth media using vented 2 L polycarbonate Erlenmeyer flasks (VWR, USA) at 27 °C. Protein expression was induced with 500 μM final concentration of sterile-filtered CuSO4. Cultured media were harvested 6 days post-induction, clarified by centrifugation at 6750×g for 20 minutes, concentrated and buffer exchanged into Ni-NTA binding buffer (20 mM Tris-HCl [pH 8.0], 300 mM NaCl, 20 mM imidazole) using a Centramate tangential flow filtration system (Pall Corp., USA) All JUNO proteins were purified by Ni-NTA metal affinity chromatography. Eluted sample was thrombin (EMD Millipore, USA) digested at 22 °C for 24 hours (1U per mg of protein) during dialysis in 1X PBS. Finally, JUNO was purified by size exclusion chromatography on a Superdex-200 Increase 10/300 column equilibrated with PBS or HBS. Peak fractions were pooled and protein concentrations were quantified by measuring A280.

### 3. Conjugation of Magnetic Beads

The conjugation of recombinant protein to the magnetic beads was performed following the modified protocol previously described in Hamze et al. (Hamze *et al*., 2020a, Hamze *et al*., 2020c). Briefly, 40 µL of magnetic sepharose beads (His Mag Sepharose Excel, GE Healthcare, Sweden) were size-selected (>70 µm) using EASYstrainer™ 70 µm filter (Greiner Bio-One, Austria) and washed in an imidazole-containing buffer to ensure the absence of any bound proteins. The washed beads were then incubated overnight at 4 °C with either 3µg of recombinant His-tagged JUNO protein (BJUNO) or with untagged recombinant JUNO protein as a control (BControl) under orbital agitation. Following incubation, the conjugated beads were washed with sodium phosphate buffer and stored at 4 °C until use.

### 4. Western Blotting

SDS-PAGE was employed to confirm the successful conjugation of recombinant proteins to magnetic beads. The beads were incubated under reducing conditions with 4X SDS Sample Buffer (Millipore, USA) for 10 min at 100 °C. Subsequently, the samples were separated by SDS-PAGE and transferred onto PVDF membrane (Immobilon-P, Millipore, USA). After incubating overnight at 4 °C with mouse anti-His antibody (1:1,000 v/v; Qiagen, Germany), the membrane was incubated for 1h at RT with HRP-conjugated anti-mouse secondary antibody (1:10,000 v/v; Thermo Fisher Scientific, USA). Immunoreactivity was revealed by chemiluminescence detection (Pierce ECL-Plus, Thermo Fisher Scientific, USA). The details of all antibodies and kits used can be found in Supplementary Table S1.

### 5. Semen Sample Preparation

#### 5.1. Fresh Samples

Three donors were recruited from Next Fertility Murcia clinic (Spain). Semen samples were obtained by masturbation following 3-5 days of sexual abstinence. After liquefaction at room temperature, the semen samples were washed in G-MOPS™ medium (1:1, v/v) and resuspended in a final volume of 2mL of G-MOPS™ (Vitrolife, Sweden). Semen parameters were determined following WHO guidelines (World Health Organization, 2021). Only samples that fulfilled the WHO reference values were included in the analysis (Supplementary Table S2).

#### 5.2. Frozen Samples

Two semen freezing methods, conventional cryopreservation and vitrification, were compared in this study. Semen samples from three different donors were used, with each ejaculate being split and frozen using both techniques. The frozen samples were provided by BiokiBank (Spain), which performed the cryopreservation and vitrification procedures. For cryopreservation, semen samples were mixed with the cryoprotectant SpermFreeze™ SSP (FERTIPRO, Belgium) at a 1:3 (v/v) ratio. The samples were then slowly frozen in straws following the standard laboratory protocol. For vitrification, semen samples were processed using the patented VitriStraw® technology (WO 2012/028967, BIOKIBANK). Briefly, spermatozoa were prepared by density gradient centrifugation, the vitrification medium was added, and the samples were subjected to ultra-rapid cooling in liquid nitrogen.

Thawing of spermatozoa was performed according to the supplier’s recommendation. For vitrified samples, 4 mL of pre-heated (42 °C) G-MOPS™ PLUS medium (Vitrolife, Sweden) was used, and the sample was left at 42 °C for 5 minutes before further processing. For the cryopreservation method, the straws were thawed in a water bath at 40 °C for 5 minutes, and the content was then transferred then into 4 mL of G-MOPS™ PLUS medium (Vitrolife, Sweden).

Motile spermatozoa were selected from both fresh and frozen samples using the swim-up method for 1 hour at 37 °C in supplemented G-MOPS™ PLUS medium (20 mM CaCl2 and 10 mg/mL progesterone). Only samples with more than 50% total motility after swim-up selection were included in the study. Motility was assessed using a computer-assisted sperm analysis (CASA) system (ISAS®, Proiser, Spain) at 37 °C. A minimum of 5 fields and 200 spermatozoa per sample were analyzed (52 frames/s, 10x magnification and Spermtrack20 chamber). Selected spermatozoa were further capacitated for 3 additional hours under the same conditions before use.

### 6. Sperm Binding Assay and Acrosome State Assessment

Groups of 30-35 magnetic sepharose beads were washed in equilibrated G-IVF™ PLUS media (37 °C, 6% CO2 in air with maximum humidity; Vitrolife, Sweden) and placed in 4-well dishes (Nunc, Denmark) containing up to 500 µL of the same medium. Capacitated spermatozoa (4-hours incubation) were added to each well at a final concentration of 200,000 or 400,000 sperm/mL. The beads and the spermatozoa were coincubated overnight at 37 °C under 6% CO2 in air at 100% humidity.

Following overnight coincubation with the beads, spermatozoa were stained with anti-CD46-FITC antibody M177 clone (3 µg/mL; Santa Cruz Biotechnology, USA) for 30 minutes at 37 °C. Afterwards, the unbound spermatozoa were collected for flow cytometry (detailed below), and the spermatozoa bound to the bead (sperm-bead complexes) were fixed with 2% paraformaldehyde and counterstained with Hoechst 33342 (1 µg/mL; Sigma-Aldrich, USA) for 30 minutes at room temperature. The fixed sperm-bead complexes were then transferred into a new well containing 0.5% (v/v) paraformaldehyde solution and evaluated within 2 hours.

The percentage of beads with at least one sperm bound, the mean number of sperm bound to each bead and its acrosome state (percentage of acrosome-reacted sperm among total bound sperm) were quantified using widefield fluorescence microscopy (Widefield Leica Thunder-TIRF imager, Germany). A multidimensional acquisition approach was applied, including multichannel imaging, z-stack analysis, multiple stage positions, and mosaic imaging. Fluorescent probes (FITC and Hoechst) were excited at 479 nm and 391 nm, respectively, with emissions detected at 519 nm and 435 nm, respectively. Acrosome-reacted cells exhibited green and blue fluorescence (CD46+ and Hoechst+), while cells with an intact acrosome displayed only blue fluorescence (CD46– and Hoechst+).

### 7. Monoclonal Antibodies inhibition assay

To evaluate the specificity of sperm binding to BJUNO via the IZUMO1–JUNO interaction, two monoclonal anti-IZUMO1 antibodies targeting two noncompeting epitopes were used (Tang *et al*., 2022). Antibody 6F02 binds to the IZUMO1 epitope involved in JUNO recognition, thereby blocking the IZUMO1–JUNO interaction. In contrast, antibody 4E04 recognizes a different epitope not implicated in JUNO binding. Murine IgG1 was included as an isotype negative control. Antibodies were added directly into the wells immediately prior to sperm insemination, at two final concentrations: 2.5⍰µg/mL and 5⍰µg/mL.

### 8. Flow Cytometry

Acrosome integrity of in-suspension spermatozoa was assessed by flow cytometry just after thawing, after swim-up selection method, and before (4h in capacitation conditions) and after co-incubation with the beads.

An aliquot of spermatozoa was mixed with Cell Staining Buffer (BioLegend, USA) and incubated with anti-CD46-FITC antibody M177 clone (3 µg/mL; Santa Cruz Biotechnology, USA) for 30 minutes in the dark before evaluation.

Fluorescent probe (FITC) was excited with a 488-nm blue solid-state laser, and its emission was detected at 530 nm, using a BD FACSCanto flow cytometer (BD Biosciences, USA). A total of 10,000 events were evaluated for each sample. Data acquisition and analysis were performed using BD FACSDiva software (BD Biosciences, USA). Acrosome-reacted cells exhibited green fluorescence (CD46+).

### 9. DNA integrity

DNA integrity was evaluated with the terminal deoxynucleotidyl transferase dUTP nick end labelling assay (TUNEL) using the In Situ Cell Death Detection Kit (Roche, Switzerland) according to Peris-Frau et al. (Peris-Frau *et al*., 2023) for unbound spermatozoa. For bead-bound spermatozoa, some modifications were introduced. Briefly, following CD46-FITC staining and 2% (v/v) paraformaldehyde fixation, the beads were permeabilized with 0.1% (v/v) Triton X-100 in PBS for 5 min and washed with PBS. DNA fragmentation was detected by incubating each group of 30 beads with 15 μL of terminal deoxynucleotidyl transferase (TdT)– fluorescent-labeled (Tetramethylrhodamine, TMR Red) nucleotide mix for 1 h in a dark, humidified chamber at 37 °C. Then washed with PBS and then counterstained with 0.1 mg/mL of Hoechst 33342 for 5 min to visualize total DNA. The sperm-bead complexes and the unbound spermatozoa were counted using a widefield fluorescence microscopy (Widefield Leica Thunderxf-TIRF imager, Germany). Fluorescent TMR Red probe was excited at 554 nm and detected at 594 nm, while FITC probe was excited at 450 nm and detected at 510 nm. Cells with fragmented DNA exhibited red fluorescence (TUNEL+), in addition to blue fluorescence (Hoechst+). In contrast, cells with intact DNA (TUNEL–) displayed only blue fluorescence (Hoechst+). Acrosome-reacted cells exhibited green and blue fluorescence (CD46+ and Hoechst+).

### 10. Statistical analysis

Data were analyzed using RStudio (R version 4.1.3) (RStudio Team, 2020). For the, presented as mean ± SEM, a generalized linear mixed-effects model (GLMM) for negative binomial family was applied given the distribution of the data. Pairwise contrasts were performed to compare groups using the Benjamini-Hochberg procedure (False Discovery Rate, FDR).

For the results presented as percentages (beads with at least one sperm bound, acrosome reaction and DNA integrity), a logistic regression for binomial data (mixed-effects model, GLMM) was used to evaluate the independence between variables. Pairwise comparisons of proportions were conducted with FDR correction.

For the classification of semen donors, an unsupervised cluster analysis was performed using the *k-means* algorithm. Donors were clustered into two groups (*k* = 2) based on the mean number of sperm bound per bead across replicates and the t-score (mean/SEM) for each donor, after standardizing and reweighting the variables by multiplying them by 1 and 0.5, respectively. Subsequently, a receiver operating characteristic (ROC) curve analysis was conducted to determine the optimal cut-off value.

## Results

### Human fresh spermatozoa bind specifically to JUNO-coated beads

Recombinant human JUNO (rhJUNO) protein was successfully conjugated to the sepharose beads. A uniform coating on the JUNO-beads surface was observed by widefield fluorescence microscopy (Figure 1A). The expected molecular weight of rhJUNO was detected both before and after conjugation to the beads, across three different protein concentrations (Figure 1B). To evaluate the stability of the JUNO protein conjugated to the beads under *in vitro* fertilization (IVF) assay conditions, JUNO-beads were incubated overnight at 37 °C, 6% CO2 in air with maximum humidity. Two different IVF media commonly used in assisted reproduction technologies were tested: Multipurpose Handling Medium-Complete (MHM-C, FUJIFILM Irvine Scientific, Inc., USA), and G-IVF™ PLUS (Vitrolife, Sweden) (Figure 1C). This assay demonstrated that the rhJUNO protein remains stably conjugated to the beads under standard IVF incubation conditions.

**Figure 1.**
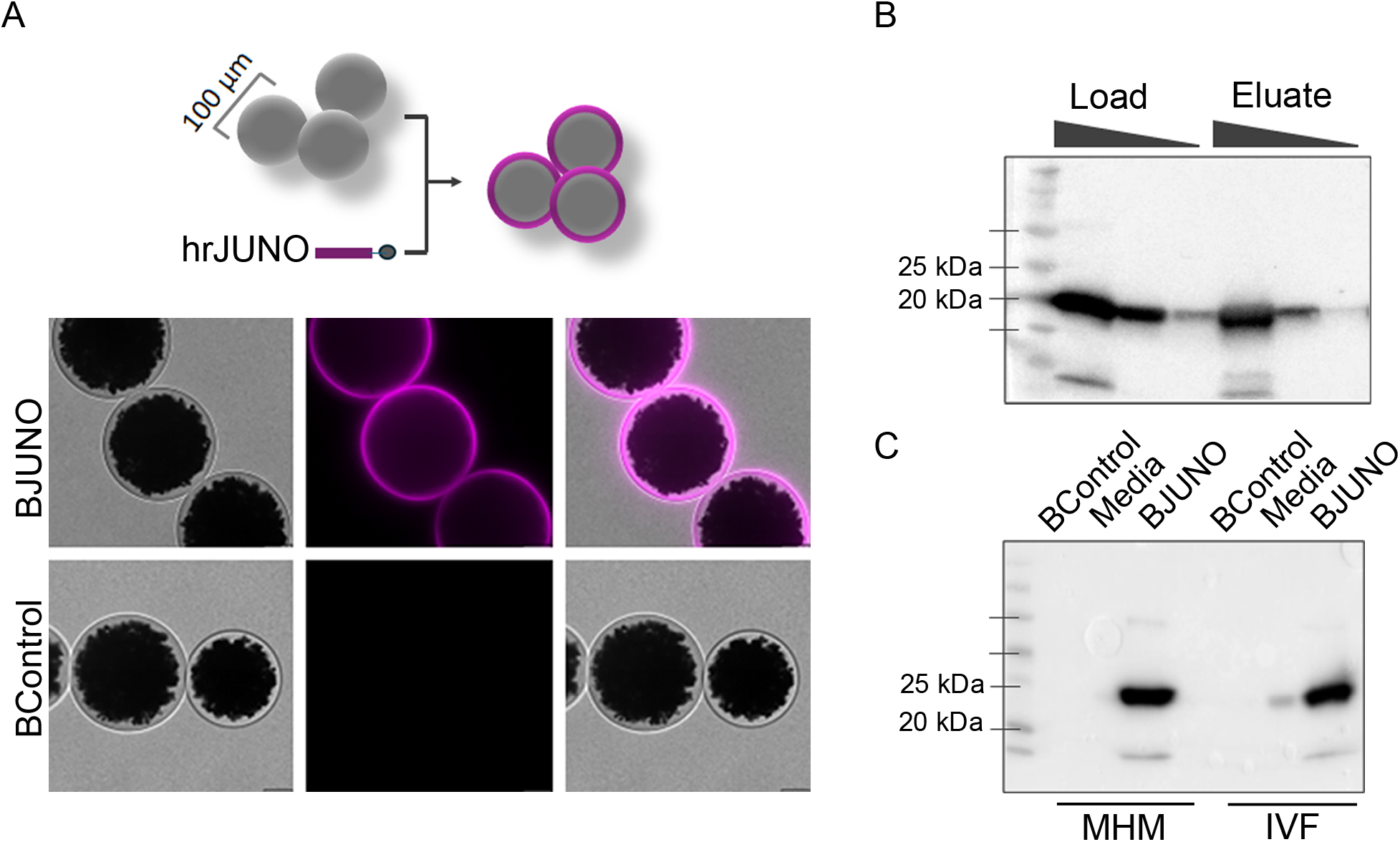
Recombinant human JUNO (rhJUNO) protein stable conjugated to beads. **A**. Schematic representation and widefield fluorescence images of BJUNO and BControl groups showing uniform rhJUNO coating on the bead surface. Scale bar: 20 µm. **B**. Different rhJUNO protein concentrations for beads conjugation were evaluated by Western Blot (upper WB). *Load* lanes indicate the decreasing amounts of rhJUNO protein tested for conjugation (3, 0.3 and 0.03 µg, respectively; ≈ 20 kDa). *Eluate* lanes represent the eluted fraction from 60 beads conjugated with each decreasing amounts of rhJUNO protein, confirming the graded reduction in protein content in the beads. **C**. Protein conjugation stability after overnight (ON) incubation was assessed using two different medias (lower WB). *BControl* and *BJUNO* lanes represent the eluates from 60 beads of each group, and the *media* lanes correspond to the BJUNO supernatant after ON incubation using MHM-C (lanes 1-3) or IVF-PLUS (lanes 4-6). No JUNO protein was detected in BJUNO supernatants from either medium, showing stable conjugation after overnight incubation at 37°C.

To validate human sperm-binding to JUNO-beads, freshly ejaculated sperm samples from three different donors were co-incubated with JUNO-beads (BJUNO) and Control-beads (BControl) at two sperm concentrations comparable to those used in IVF protocols. After the incubation period commonly used in IVF procedures, sperm bound to the beads (identified as Hoechst-positive blue dots) (Figure 2A) were quantified to evaluate two key parameters of binding performance for each sample: (i) the average number of sperm cells attached per bead, and (ii) the proportion of beads with at least one attached sperm cell, as a measure of binding specificity. The evaluation under a fluorescent microscope showed that BJUNO group showed a significantly higher number of bound sperm cells compared to the BControl group (p ≤ 0.0001). Remarkably, increasing the concentration led to a 150% rise in the number of attached sperm cells in BJUNO, from 4.4 ± 0.5 to 6.4 ± 0.8 (*p* < 0.001). In contrast, this increase had no significant effect on the control group, where the number of bound sperm cells remained relatively unchanged (0.6 ± 0.1 and 0.9 ± 0.1, respectively; p = 0.056) (Figure 2B). When evaluating the percentage of beads with at least one bound sperm at both sperm concentrations, we observed that the majority of BJUNO had sperm attached (88.3%, CI 95%: 81.4% – 93.3%, n = 128), whereas in the control group, most beads showed no bound sperm (43.2%, CI 95%: 33.9% – 53.0%, n = 111) (*p* < 0.0001) (Figure 2C). Taken together, these results indicate that human sperm bind specifically and, in a dose-dependent manner to BJUNO, and that this binding capacity to the beads model is dependent on the presence of the receptor, JUNO.

**Figure 2.**
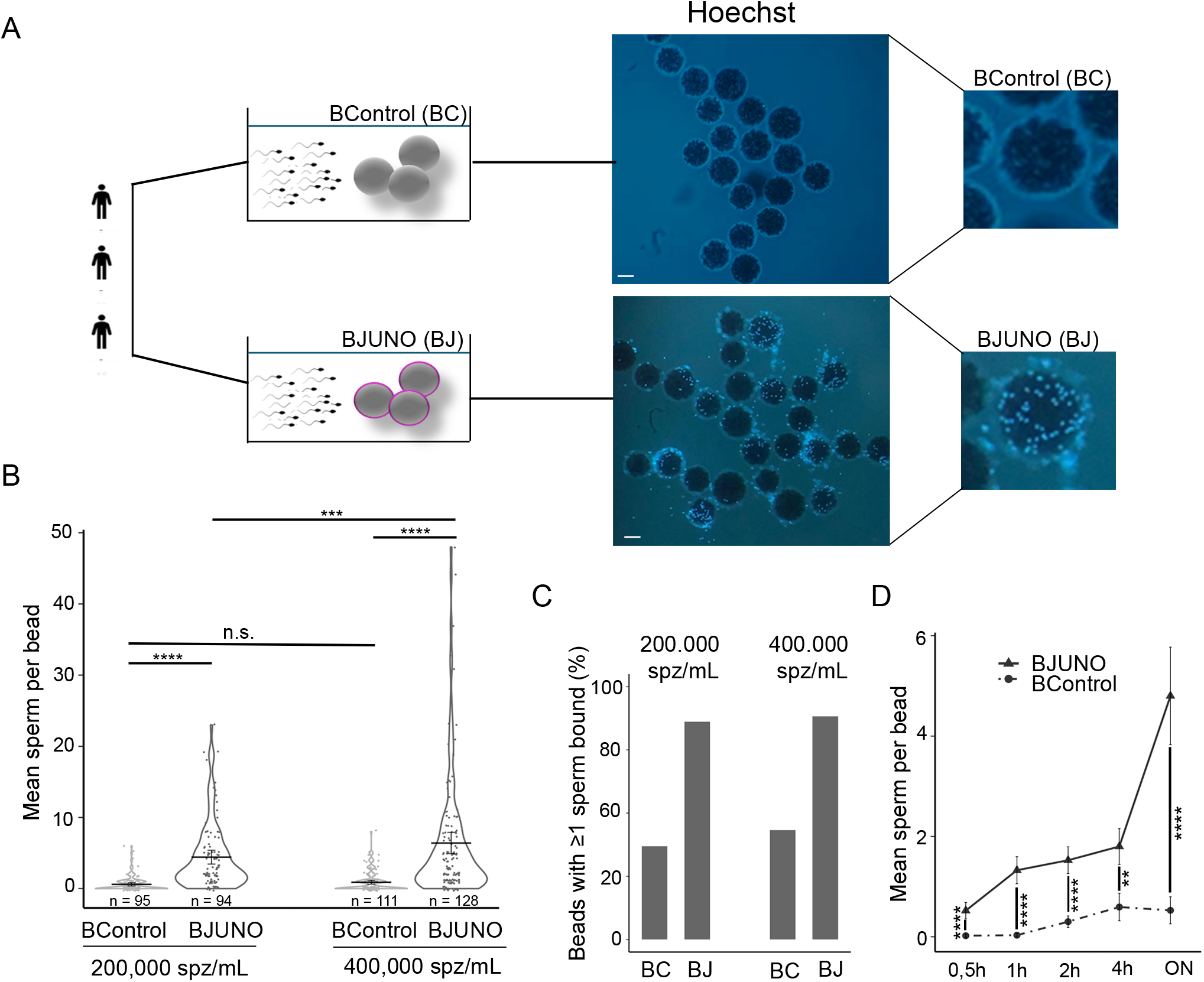
Sperm binding assay to BJUNO and BControl using human fresh semen samples. **A**. Schematic and epifluorescence microscopy images (Hoechst) of BControl (BC) and BJUNO (BJ) after co-incubation with spermatozoa from three different donors. Scale bar: 100 µm. *B*. Quantification of sperm-bead interaction under two sperm concentrations (200,000 and 400,000 spermatozoa/mL). Only spermatozoa whose heads were in contact with the bead were counted. **B**. Mean number of spermatozoa per bead (violin plot). **C**. Percentage of beads with at least one sperm bound (bar plot). Both parameters were significantly higher for BJUNO compared to BControl. **D**. Time-course analysis (line plot) of sperm binding dynamics, showing a progressive increase in sperm attachment to BJUNO over time, with overnight (ON) incubation yielding the highest number of sperm per bead. Abbreviations: n.s., not significant; \**p* < 0.05; \*\**p* ≤ 0.01; \*\*\**p* ≤ 0.001; \*\*\*\**p* ≤ 0.0001.

We subsequently selected a dose of 400,000 spermatozoa/mL to maximize the number of bound sperm and, consequently, increase the sensitivity of the assay. To determine the optimal incubation time, binding kinetics were assessed at 30 min, 1 h, 2 h, 4 h, and overnight. Sperm binding to BJUNO increased progressively, with significant differences from BControl evident as early as 30 min (p < 0.0001). The greatest difference was observed after overnight incubation, with an average of 4.8 ± 0.5 sperm bound per bead in the BJUNO group (n = 90) versus 0.5 ± 0.1 in the BControl group (n = 91) (*p* < 0.0001) (Figure. 2D; Supplementary Table S3). A similar trend was observed when analysing the percentage of beads with at least one bound sperm (Supplementary Figure, Table S3). These results indicate that the specific binding of human sperm to BJUNO is also time dependent.

### Vitrified human sperm bind specifically to JUNO beads

To validate the model, we compared the binding capacity of sperm samples subjected to two different cryopreserved methods, based on the premise that such procedures can affect membrane stability (Barthelemy *et al*., 1990) and, consequently, the presentation of receptors involved in sperm-oocyte binding. For this analysis, ejaculates from three donors were split into two aliquots: one processed using a Cryopreserved-based method, while the other was frozen by vitrification (Figure. 3A, Supplementary Table S4). After thawing, samples were incubated with BJUNO and BControl and analysed as previously described.

**Figure 3.**
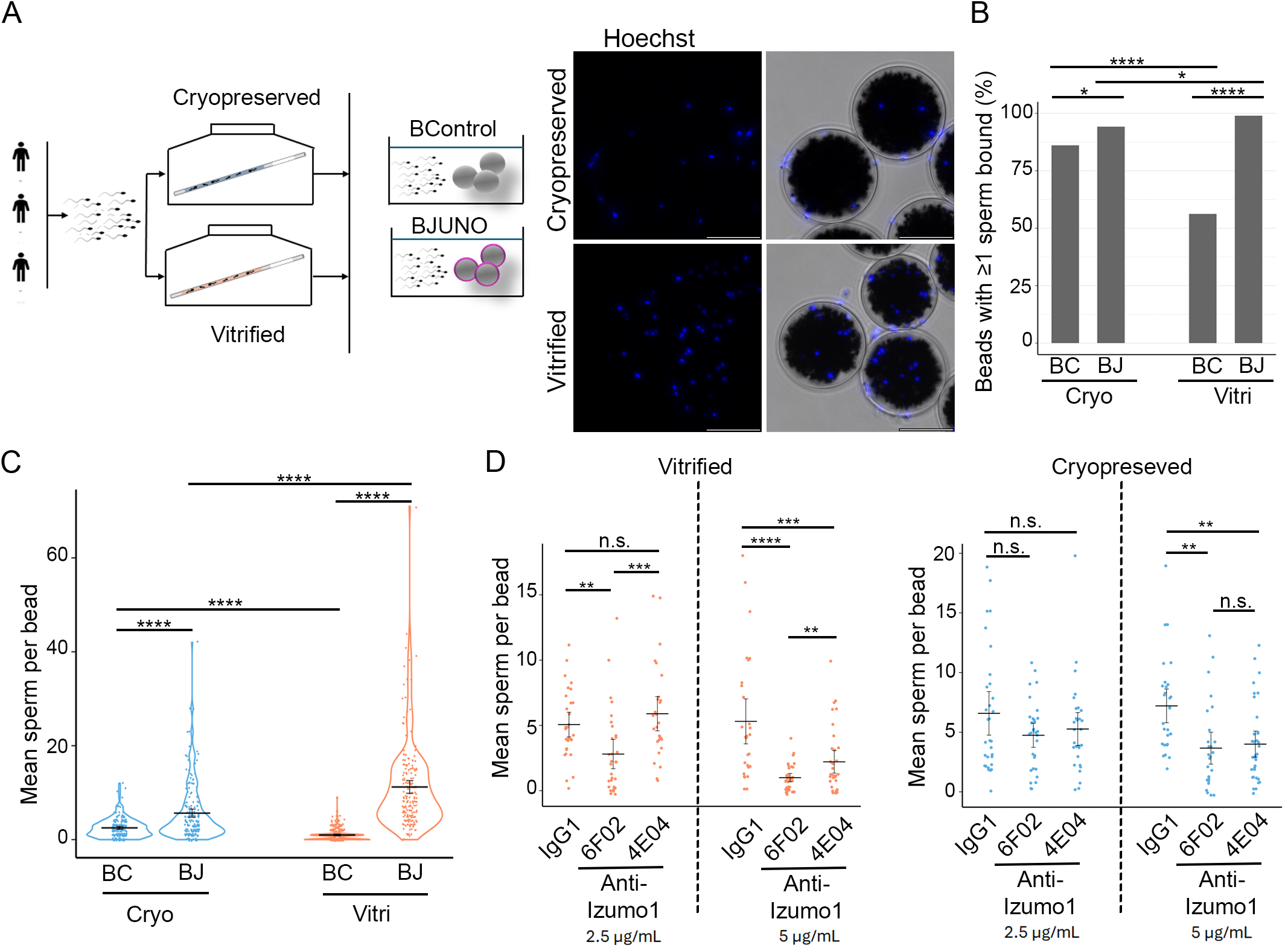
Effect of freezing method on sperm binding to JUNO-beads. **A**. Schematic representation of the experimental groups (left panel) and the widefield fluorescence images (Hoechst) of the sperm binding assay (right panel). Only spermatozoa with their heads in contact with the bead were counted. Scale bar: 100 µm. **B**. The percentage of beads with at least one sperm bound was quantified for both cryopreserved and vitrified samples. BJUNO (BJ) showed a consistently higher binding compared to BControl (BC) in both types of preserved samples. **C**. The mean number of spermatozoa bound per bead was compared between BJ and BC for each preservation method. A significantly higher binding was observed for BJUNO, with the difference between BJ and BC being more pronounced in vitrified samples. **D**. Specificity of the sperm binding assay was tested using anti-IZUMO1 antibodies (4E04 and 6F02) at two concentrations (2.5 µg/mL, left; and 5 µg/mL, right). These antibodies target different epitopes of IZUMO1, with 6F02 specifically blocking the epitope involved in JUNO binding. IgG1 was used as a negative antibody control. In cryopreserved samples, no significant difference was observed at 2.5 µg/mL, while both antibodies reduced sperm binding at 5 µg/mL. In vitrified samples, 6F02 caused a clear reduction in sperm binding at both concentrations, with stronger inhibition at 5µg/mL. Abbreviations: n.s., not significant; \**p* < 0.05; \*\**p* ≤ 0.01; \*\*\**p* ≤ 0.001; \*\*\*\**p* ≤ 0.0001.

We observed that vitrified samples exhibited a significantly higher proportion of BJUNO with at least one bound sperm (98.9%, 95% CI: 95.8%–99.7%) compared to BControl (56.3%, 95% CI: 49.0%–63.3%; p ≤ 0.0001), consistent with the results obtained from fresh samples. In cryopreserved samples, although a statistically significant difference was still observed between BJUNO (94.2%, 95% CI: 89.9%–96.8%) and BControl (86.0%, 95% CI: 80.3%–90.3%; *p* < 0.05), the magnitude of the difference was smaller. Moreover, in the cryopreserved group, a larger proportion of BControl presented at least one bound sperm, suggesting a higher number of sperm undergoing nonspecific binding (Figure 3B).

Regarding the average number of sperm bound per bead (Figure 3C), vitrified samples again showed superior binding efficiency, with BJUNO displaying 11.2 ± 0.7 bound spermatozoa (n = 188) compared to 0.98 ± 0.1 in BControl (n = 183). In Cryopreserved samples, binding was lower overall, with 5.6 ± 0.4 sperm bound to BJUNO (n = 191) and 2.5 ± 0.2 to BControl (n = 186). While both methods demonstrated preferential binding to BJUNO over BControl (*p* ≤ 0.0001), sperm from vitrified samples exhibited significantly greater binding efficiency compare with sperm from slow freezing (p ≤ 0.0001). Additionally, the number of sperm bound to BControl was significantly lower in vitrified samples compared to cryopreserved ones (*p* ≤ 0.0001), further supporting that the vitrified method better preserves the cellular and molecular conditions necessary for specific binding to BJUNO.

To demonstrate that human sperm binding to BJUNO is specifically mediated by the IZUMO1– JUNO interaction, two well-characterized anti-IZUMO1 antibodies targeting distinct epitopes were used (Figure 3D). Anti-IZUMO1 6F02 blocks the epitope directly involved in IZUMO1-JUNO binding, whereas anti-IZUMO1 4E04 recognizes a different region not implicated in this interaction (Tang *et al*., 2022). In vitrified samples (Figure 3D, left graph), the use of anti-IZUMO1 6F02 at 2.5⍰µg/mL significantly reduced sperm binding compared to the control (2.8 ± 0.6 vs. 5.1 ± 0.5; *p* < 0.01), confirming specific inhibition of the IZUMO1-mediated interaction. In contrast, anti-IZUMO1 4E04 had no significant effect at the same concentration (5.9 ± 0.7; *p* > 0.05). When the antibody concentration was increased to 5⍰µg/mL, both antibodies reduced sperm binding, but the effect was significantly stronger with anti-IZUMO1 6F02 (1.0 ± 0.2 vs. 5.3 ± 0.9; *p* < 0.0001).

In cryopreserved samples (Figure 3D, right graph), the inhibitory effect of the antibodies was less pronounced. At 2.5⍰µg/mL, neither anti-IZUMO1 6F02 nor 4E04 significantly reduced sperm binding compared to the control (4.7 ± 0.5, 5.3 ± 0.7, and 6.6 ± 0.9, respectively; *p* > 0.05). At 5⍰µg/mL, both antibodies caused a moderate but comparable reduction (3.7 ± 0.7, 4.0 ± 0.6, and 7.2 ± 0.7, respectively; *p* < 0.01), suggesting that in these samples, in addition to the specific IZUMO1-JUNO interaction, nonspecific binding mechanisms may also contribute.

A key cellular requirement for gamete interaction is that spermatozoa must be acrosome-reacted. Loss of the acrosome leads to the localization of IZUMO1 at the equatorial segment of the sperm head, enabling its interaction with JUNO present on the oocyte membrane (Satouh *et al*., 2012; Inoue and Wada, 2018; Manjon *et al*., 2025). Therefore, sperm that bind specifically to JUNO-coated beads are expected to be acrosome-reacted. To investigate this, we first assessed the acrosomal status of cryopreserved and vitrified sperm samples under different incubation times and preparation protocols (Supplementary Figure S1), using CD46 as a marker for acrosome-reacted live spermatozoa (CD46^+^). The most significant difference was observed after overnight incubation, where vitrified samples showed a significantly higher percentage of acrosome-reacted cells (19.2%) compared to cryopreserved samples (10.0%) (*p* < 0.0001), as determined by flow cytometry. After establishing the acrosomal status under our experimental conditions, we then evaluated the acrosomal status of spermatozoa in the sperm–bead binding assay after overnight incubation, comparing sperm bound to the beads versus those remaining unbound. In the unbound population (Figure 4A), the proportion of acrosome-reacted spermatozoa was relatively low (29.1% ± 11.8 in cryopreserved and 41.7% ± 13.8 in vitrified samples (Figure 4A, Unbound sperm graph). In contrast, most of sperm bound to the beads were acrosome-reacted, with over 85% being CD46^+^ in both conditions (86.4% ± 1.6 for cryopreserved and 96.7% ± 0.7 for vitrified samples) (Figure 4A, Bound sperm graph), indicating that binding to JUNO-coated beads selectively occurs in acrosome-reacted spermatozoa.

**Figure 4.**
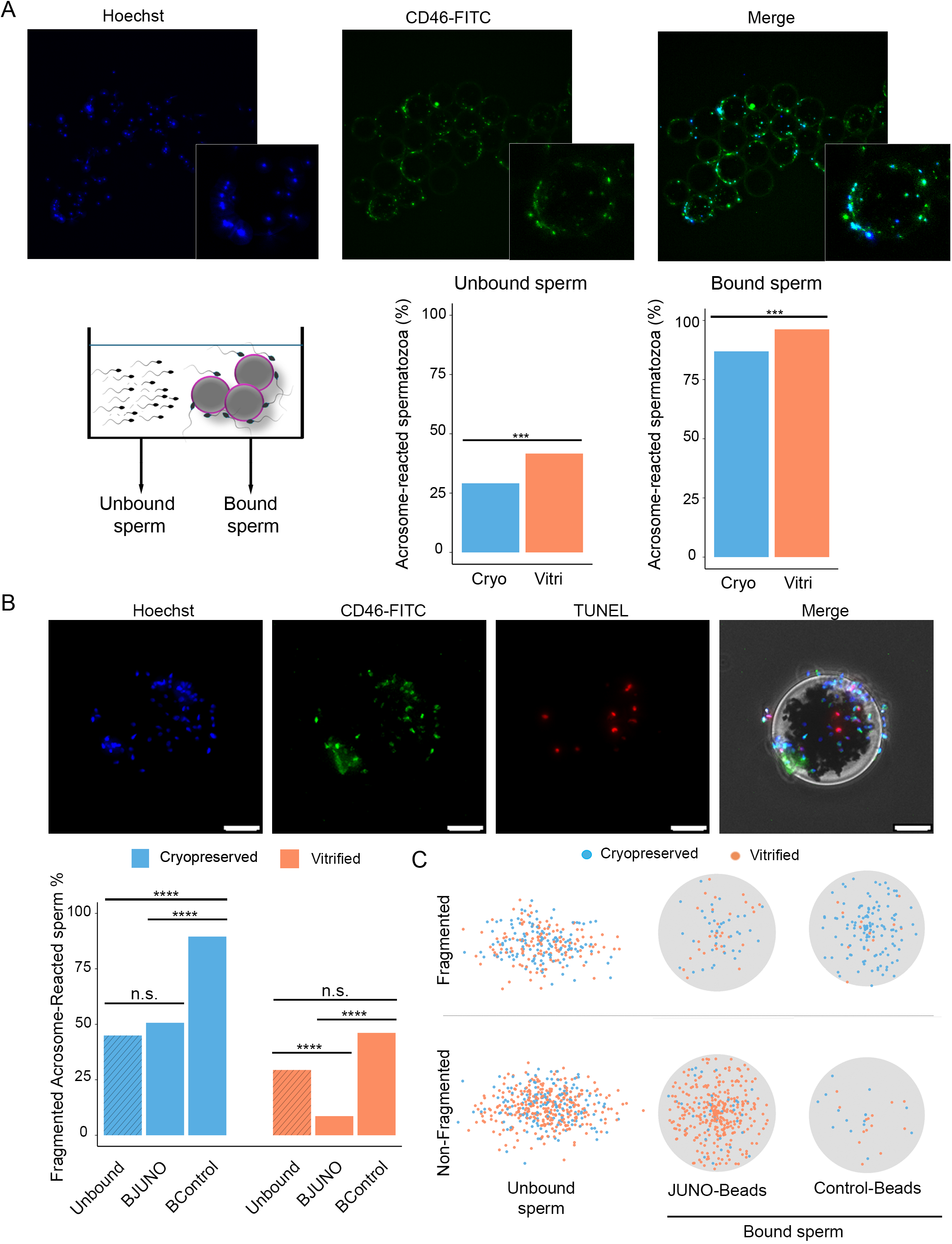
Acrosome reaction and DNA fragmentation evaluation on sperm-JUNO bead binding assay. **A**. The percentage of acrosome-reacted spermatozoa (Hoechst+ and CD46+) was quantified in unbound sperm by flow cytometry (left bar plot) and BJUNO-bound sperms by widefield fluorescence (right bar plot), revealing a ∼50% enrichment of acrosome-reacted spermatozoa in the BJUNO group across both types of preserved samples. **B**. DNA integrity of acrosome-reacted spermatozoa was assessed by TUNEL assay (widefield fluorescence) in both bead-bound and unbound cells (upper panels). Scale bar: 10 µm. The percentage of acrosome-reacted spermatozoa with DNA fragmentation (left bar plot) was higher in cryopreserved samples than in vitrified ones. Additionally, BJUNO-bound spermatozoa showed lower levels of DNA fragmentation compared to BControl-bound and unbound sperm. *C*. The individual cell distribution according to DNA integrity of acrosome-reacted spermatozoa is represented in the right dot plot. Abbreviations: n.s., not significant; \**p* < 0.05; \*\**p* ≤ 0.01; \*\*\**p* ≤ 0.001; \*\*\*\**p* ≤ 0.0001.

Acrosome reaction could be associated with, and may be both a cause and consequence of, sperm cell death (Harper *et al*., 2008). To determine whether sperm bound to JUNO-coated beads undergo acrosome reaction as a functional requirement for receptor interaction, rather than as a degenerative event, we evaluated DNA integrity in acrosome-reacted sperm using the TUNEL assay (Figure 4B). In vitrified samples, 29.4% of unbound reacted-spermatozoa exhibited DNA fragmentation. In contrast, reacted-spermatozoa bound to JUNO beads showed a significantly lower rate of DNA fragmentation (<10%; *p* < 0.0001) (Figure 4B, orange graph) indicating that most JUNO-bound sperm are acrosome-reacted and maintain intact DNA, a hallmark of functional viability. This contrasts with the BControl-bound group, in which 46.2% of acrosome-reacted spermatozoa exhibited DNA fragmentation (*p* > 0.05 vs. unbound), suggesting that damaged sperm may bind non-specifically (Figure 4B, orange graph). A similar pattern was observed in cryopreserved semen samples (Figure 4B, blue graph), where 45.0% of unbound spermatozoa showed DNA fragmentation, and a comparable percentage was found among BJUNO-bound spermatozoa (50.6%; *p* > 0.05). In contrast, the Control-bound fraction displayed significantly higher DNA fragmentation (89.5%; *p* < 0.0001 vs. unbound) (Figure 4B, blue graph). These results suggest that cryopreserved samples bind more non-specifically to surfaces, likely due to increased cellular damage.

When acrosome-reacted spermatozoa were further classified based on the presence or absence of DNA fragmentation, represented by individual sperm counts (dots) across assay conditions: unbound, JUNO-beads, and control beads, a clear pattern emerged (Figure 4C). The population of non-fragmented, acrosome-reacted spermatozoa was most enriched among vitrified samples bound to JUNO-beads, whereas in cryopreserved samples, a greater number of DNA fragmented spermatozoa were bound to control beads. These findings indicate that cryopreserved samples not only show reduced binding to JUNO-beads, but also exhibit increased non-specific binding, likely as a result of greater cellular damage. This is supported by the observed DNA fragmentation, which may be linked to membrane destabilization caused by the freezing process, potentially leading to mislocalization of the IZUMO1-binding complex.

Together, these results demonstrate that JUNO-coated beads selectively bind vitrified spermatozoa exhibiting features essential for fertilization, namely, acrosome-reacted status and low DNA fragmentation, underscoring the specificity and biological relevance of the assay.

### Sperm-bead binding assay as a functional evaluation of semen quality

Once we validated that the vitrified sperm bound to JUNO-beads exhibit the characteristic features of fertilization-competent sperm and that their binding can be quantitatively assessed, we proceeded to compare the binding capacity to JUNO-beads across normalized semen samples from different donors. This approach allows us to evaluate inter-individual variability and supports the potential of the sperm-beads binding assay as a functional test for sperm quality. To this end, the sperm-bead binding assay was performed using vitrified sperm from 18 individual donors (29 ejaculates and 50 replicates), analyzing over 3,300 beads. Consistent with previous observations, the mean number of spermatozoa bound to JUNO beads was significantly higher than that bound to Control beads (8.2 ± 0.2, n = 1,679 vs. 0.7 ± 0.0, n = 1,689; *p* < 0.0001) (Supplementary Figure S2; Supplementary Table S5). To classify donors based on their sperm-bead binding performance, an unsupervised clustering analysis using k-means was performed on donors with at least two replicates (n=12). Twelve donors were grouped into two categories based on two features: the mean number of sperm bound per bead and the t-score (mean/SE) calculated across replicates (Figure 5A). The k-means algorithm categorized donors according to the similarity of these features, yielding two clusters; six donors, which we classified as low binding capability (LBC), and six as high binding capability (HBC). A receiver operating characteristic (ROC) curve was subsequently generated using the mean sperm count per bead as a discriminating variable, identifying an optimal cut-off value of 9.06 spermatozoa per bead (Figure 5A). These results indicate that the sperm-bead binding assay can be used to classify individuals based on the ability of their semen samples to bind to JUNO-beads. Additional validation using independent donor samples will be necessary to evaluate the robustness and general applicability of this classification approach.

**Figure 5.**
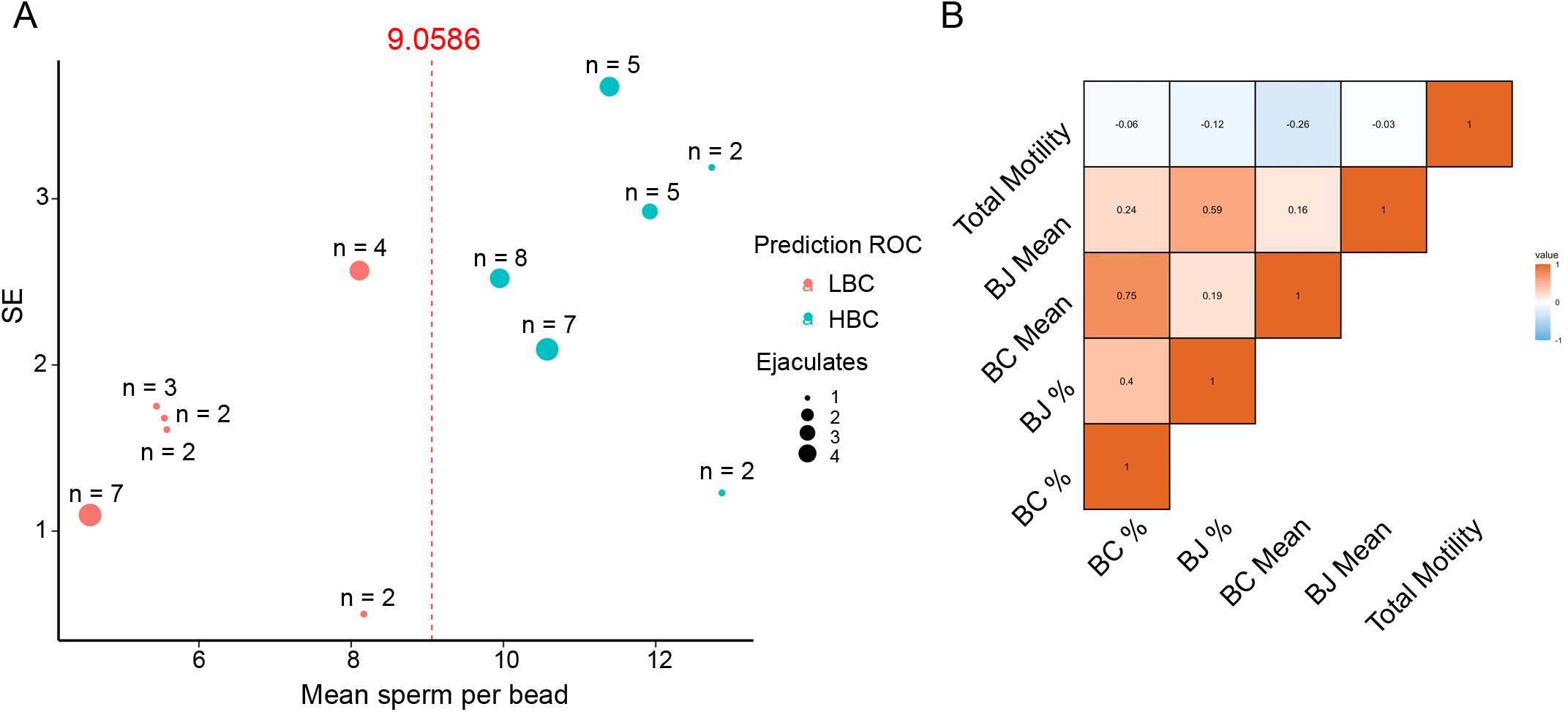
Validation of the sperm binding assay. **A**. Representation of semen donors’ classification based on the mean number of sperm bound per bead and its standard error (SE) across replicates. Dot size reflects the number of different ejaculates per donor, and the number next to each dot indicates the number of replicates. Donors classified as low binding capability (LBC) are shown in red, and those classified as high binding capability (HBC) in blue. **B**. Pearson correlation analysis between total sperm motility prior to co-incubation (Total Motility) and the two main readouts of the sperm binding assay: the mean number of spermatozoa bound per bead (BJ Mean and BC Mean) and the percentage of beads with at least one sperm bound (BJ % and BC %). Weak linear relationships were observed between total motility and the assay readouts.

Finally, we assessed whether the mean number of spermatozoa bound per bead or the percentage of beads with at least one sperm bound were linearly associated with total sperm motility prior to co-incubation (Figure 5B). Pearson correlation analyses revealed very weak associations between motility and all binding parameters (|r| < 0.27), indicating a negligible or absent linear relationship.

## Discussion

The model presented builds upon a previously described and validated system tested in other species using recombinant proteins involved in gamete interaction (Hamze *et al*., 2020a, Hamze *et al*., 2020c, Hamze *et al*., 2020b). In the present study, we adapted the 3D-bead-based model for human application using recombinant human JUNO protein. The approach is designed to mimic *in vitro* fertilization (IVF) by replacing the oocyte with JUNO-coated beads that serve as molecular decoys, enabling the selection of high-fertilizing sperm capable of specifically binding to JUNO. To establish proof of concept, the experimental conditions, such as sperm concentration, incubation time, and media, were aligned with standard human IVF protocols.

Our findings reveal a clear time- and concentration-dependent pattern of sperm binding specifically to JUNO-coated beads, consistent with the highly mechanostable multistate catch-bond mechanism recently described for the IZUMO1:JUNO interaction (Boult *et al*., 2025). This mechanism enables stable long-term adhesion between sperm and egg, and was also observed using the JUNO-bead. Both the number of spermatozoa attached per bead and the proportion of beads with at least one bound sperm were significantly higher for JUNO-beads compared to control beads. This strong signal-to-noise ratio stands in sharp contrast to findings in bovine samples using the same approach (Hamze *et al*., 2020c), where no significant differences were observed in the proportion of beads with at least one bound sperm. This species-specific disparity underscores that, under the tested conditions, human sperm exhibit a markedly higher efficiency in binding to JUNO-coated beads. These results also emphasize that both metrics, the number of sperm per bead and the proportion of beads with at least one bound sperm, are essential for a comprehensive interpretation of assay performance. Remarkably, efficient binding to JUNO-beads was achieved with substantially fewer spermatozoa (∼6,000– 7,000 per bead) compared to the typical sperm-to-oocyte ratios used in IVF (∼50,000) (Oseguera-López *et al*., 2019). More than 90% of JUNO-beads captured at least one sperm, enabling bead-by-bead evaluation within groups and greatly increasing the statistical robustness, and therefore the sensitivity, of the assay.

The preservation of gametes and embryos has been a cornerstone advance in assisted reproductive technologies, ensuring a reliable source of high-quality samples for both clinical and research applications. Vitrification of oocytes and embryos, in particular, has proven highly effective because it minimizes membrane damage and preserves cellular integrity and viability (Valojerdi and Salehnia, 2005; Rienzi *et al*., 2016). Extending this strategy to sperm, vitrification offers the potential to maintain optimal cellular conditions, thereby improving the functional competence of preserved spermatozoa.

The novelty of the present study lies in the evaluation of JUNO-binding capacity in spermatozoa subjected to different preservation protocols. Our findings reveal that vitrified sperm retain a higher capacity for specific binding to JUNO-beads compared to standard cryopreservation. In contrast, sperm samples subjected to slow-rate cryopreservation exhibited a pronounced increase in non-specific binding, indicating that vitrification preserves the specific functionality of IZUMO1 required for fertilization-competent interactions, IZUMO1:JUNO.

Recent studies have described a complex molecular architecture underlying gamete recognition, in which IZUMO1 is part of a major sperm membrane protein complex (Ellerman *et al*., 2009; Contreras *et al*., 2022; Lu *et al*., 2023; Pacak *et al*., 2023) where IZUMO1, SPACA6, and TMEM81 specifically interact to form a functional complex (Deneke *et al*., 2024). This trimeric complex interacts with the oocyte proteins JUNO and CD9, forming a hetero-pentameric structure at the fertilization synapse (Elofsson *et al*., 2024). Specific monoclonal antibodies, 4E04 and 6F02, both recognize IZUMO1 at non-competing epitopes; notably, the 6F02 epitope overlaps with the IZUMO1-binding site for JUNO (Tang *et al*., 2022). Consistent with the requirement to assemble this molecular structure to execute sperm–oocyte binding, in vitrified sperm samples we observed that binding was inhibited by 6F02 but not by 4E04, except at high concentrations where partial inhibition likely reflects steric hindrance. In contrast, this inhibitory effect was absent in cryopreserved sperm, suggesting that the molecular organization required for specific JUNO-mediated binding is disrupted under these preservation conditions.

To further characterize JUNO bead-bound sperm, we evaluated their acrosomal status and DNA integrity. These analyses demonstrated that JUNO-coated beads selectively capture vitrified spermatozoa exhibiting hallmarks of fertilization competence, specifically, acrosome-reacted cells with minimal DNA fragmentation (Santi *et al*., 2018; Tanga *et al*., 2021; Morohoshi *et al*., 2023). In contrast, although a large proportion of cryopreserved spermatozoa also underwent the acrosome reaction, a significant fraction displayed DNA fragmentation. This pattern suggests that cryopreservation and slow freezing compromise membrane integrity, leading to premature loss of acrosomal contents and triggering apoptotic pathways associated with DNA damage (Duru *et al*., 2001; Ozkavukcu *et al*., 2008).

Together, these results indicate that cryopreservation compromises the specificity of IZUMO1– JUNO interactions by inducing membrane damage (Ozkavukcu *et al*., 2008) that disrupts the spatial organization of receptors (Wang *et al*., 2014; Fukuda *et al*., 2016; Peris-Frau *et al*., 2020). This loss of structural integrity increases the contribution of non-specific interactions, in sharp contrast to vitrified sperm, which maintains a molecular architecture conducive to selective and functional binding. The application of sperm vitrification in vitro fertilization techniques may represent an improvement in sperm quality with respect to fertilizing capacity, which could in turn contribute to enhanced embryo viability and development.

Finally, we applied the bead-based model to classify semen donors according to their JUNO-binding capacity. Although this approach requires further validation, our results highlight the potential of this assay to discriminate sperm quality beyond conventional semen parameters. A more detailed analysis could reveal inter-individual differences in fertilization competence that stem from intrinsic molecular characteristics or subclinical pathologies. Interestingly, no significant correlation was found between total sperm motility prior to insemination and JUNO-binding efficiency during co-incubation. This finding contrasts with studies that associate motility with fertilization outcomes (Zinaman *et al*., 2000; World Health Organization, 2021; Villani *et al*., 2022). It is worth noting, however, that all semen samples analyzed here were obtained from healthy donors and fulfilled the inclusion criterion of >50% total motility, thus representing a relatively homogeneous population. Within this restricted range, motility appears to lose its predictive value for fertilization potential. These observations support the notion that, although motility is important, it is not sufficient on its own to predict fertilization competence, reinforcing the need for functional assays that capture additional sperm attributes (Guzick *et al*., 2001; Liu *et al*., 2004; Mariappen *et al*., 2018).

Collectively, these findings demonstrate that vitrification better preserves the structural and molecular integrity of sperm, enabling specific and functional interactions with JUNO-coated beads, while slow-rate cryopreservation induces membrane and DNA damage that favors non-specific binding. Our bead-based model therefore emerges as a sensitive tool to discriminate between preservation methods and to identify spermatozoa with true fertilization competence. And underscore the value of the JUNO-bead binding assay as a functional biomarker that goes beyond conventional semen analysis, providing a more precise assessment of the molecular competence of sperm to achieve fertilization.

## Author’s roles

P.C.R. and M.J.M. conceptualized and designed the study. P.C.R., M.S.T. and J.G.H. performed the experiments and acquired the data. E.G. coordinated patient recruitment from the clinic and advised on sample treatment protocols according to clinical standards. P.Y. and J.E.L. produced the recombinant protein used to generate JUNO-beads. P.C.R. and M.J.M. analysed and interpreted the data. M.J.M. supervised the project. P.C.R. and M.J.M. drafted the original manuscript. J.E.L. and E.G. provided critical intellectual revision of the manuscript. M.J.M. and J.E.L. secured funding. All authors reviewed and approved the final version of the manuscript.

## Acknowledgements

The authors want to thank Dr. S. Tang (University of Yale) for kindly providing the anti-IZUMO1 antibodies. They also gratefully acknowledge the Microscopy Service of the Scientific and Technical Research Area at the University of Murcia for technical support, and in particular, Dr. María Teresa Coronado for her valuable assistance and contribution to this work.

## Data availability

All data needed to evaluate the conclusions in the paper are present in the paper.

