## Supplementary Table S1 for "JUNO-Coated Beads as a Functional Assay to Capture and Characterize Fertilization-Competent Human Sperm"

**Supplementary Table S1. Summary of the antibodies and commercial kits used in the study**

| Reagents | Reference code | Source (company or reference) |
|---|---|---|
| Antibodies | | |
| Penta·His antibody, BSA free | 34660 | Qiagen |
| Donkey anti-mouse IgG (H+L) Cross Absorbed Secondary Antibody, HRP conjugate | SA1-100 | Thermo Fisher Scientific |
| Monoclonal mouse anti-IZUMO1 antibody (6F02) | 6F02 | Tang et al., 2022 |
| Monoclonal mouse anti-IZUMO1 antibody (4E04) | 4E04 | Tang et al., 2022 |
| Monoclonal mouse anti-CD46-FITC antibody (M177 clone) | sc-52647 | Santa Cruz Biotechnology |
| bisBenzimide H 33342 trihydrochloride | 14533 | Sigma-Aldrich |
| Kits | | |
| Pierce™ ECL Plus Western Blotting Substrate | 32132 | Thermo Fisher Scientific |
| In Situ Cell Death Detection Kit, TMR red | 12156792910 | Roche |
