## Supplementary Table S2 for "JUNO-Coated Beads as a Functional Assay to Capture and Characterize Fertilization-Competent Human Sperm"

**Supplementary Table S2. Semen parameters of fresh samples**

| | Semen donors | | |
|---|---|---|---|
| | M3 | M4 | M10 |
| **Baseline semen parameters** (standardized assessment in the clinic) | | | |
| Donor age (years) | 24 | 36 | 31 |
| Volume (mL) | 3.0 | 2.5 | 2.0 |
| pH | 8.5 | 8.7 | 8.7 |
| Total motility (%) | 72 | 59 | 41 |
| Progressive motility (%) | 62 | 54 | 38 |
| Normal morphology (%) | 12 | 9 | 14 |
| **Post-processing sperm parameters** (lab assessment) | | | |
| Sperm concentration after Swim-up (M spz/mL) | 26 | 48 | 5 |
| Total motility after Swim-up (%) | 72 | 85 | 35 |
| Sperm concentration before co-incubation (M spz/mL) | 28 | 29 | 6 |
| Total motility before co-incubation (%) | 72 | 66 | 46 |
