## Supplementary Table S3 for "JUNO-Coated Beads as a Functional Assay to Capture and Characterize Fertilization-Competent Human Sperm"

**Supplementary Table S3. Key parameters of binding performance at different co-incubation times, including mean number of spermatozoa bound to the bead and the percentage of bead with at least one sperm bound.**

| Co-incubation time | BControl | BJUNO | *p* value |
|---|---|---|---|
| | Mean ± SE (n) | Mean ± SE (n) | |
| 30 minutes | 0.02 ± 0.02 (91) | 0.5 ± 0.1 (94) | < 0.0001 |
| 1 hour | 0.03 ± 0.02 (93) | 1.3 ± 0.1 (98) | < 0.0001 |
| 2 hours | 0.3 ± 0.06 (93) | 1.5 ± 0.1 (97) | < 0.0001 |
| 4 hours | 0.6 ± 0.1 (91) | 1.8 ± 0.2 (95) | 0.0017 |
| Overnight | 0.5 ± 0.1 (91) | 4.8 ± 0.5 (90) | < 0.0001 |
| | Percentage (CI 95%) | Percentage (CI 95%) | |
| 30 minutes | 2.2 (0.4 – 8.5) | 36.2 (26.7 – 46.8) | < 0.0001 |
| 1 hour | 3.2 (0.8 – 9.8) | 65.3 (54.9 – 74.5) | < 0.0001 |
| 2 hours | 25.8 (17.5 – 36.1) | 74.2 (64.2 – 82.3) | < 0.0001 |
| 4 hours | 31.9 (22.7 – 42.6) | 73.7 (63.5 – 81.9) | 0.0028 |
| Overnight | 25.3 (17.0 – 35.7) | 91.1 (82.7 – 95.8) | < 0.0001 |

Abbreviations: BControl, beads control; BJUNO, beads JUNO; SE, standard error; CI, confidence interval.
