## Supplementary Table S4 for "JUNO-Coated Beads as a Functional Assay to Capture and Characterize Fertilization-Competent Human Sperm"

**Supplementary Table S4. Donor characteristics and paied sperm parameters of the same samples preserved by cryopreservation and vitrification**

| | Semen donors | | | | | |
|---|---|---|---|---|---|---|
| | D045 | | D089 | | D048 | |
| **Biological Donor Parameters** | | | | | | |
| Age (years) | 42 | | 19 | | 22 | |
| Blood group (ABO and Rh) | O- | | A+ | | A+ | |
| Proven fertility | Yes | | Yes | | Yes | |
| **Freezing method** | Cryo | Vitri | Cryo | Vitri | Cryo | Vitri |
| **Post-thawing semen characteristics according to biobank** | | | | | | |
| Progressive motility (%) | 40 | 78 | 42 | 86 | 33 | 77 |
| Normal morphology (%) | 9 | 9 | 6 | 6 | 8 | 8 |
| **Post-thawing semen characteristics (laboratory-assessed)** | | | | | | |
| Sperm concentration after thawing (M spz/mL) | 34 | 7 | 15 | 5 | 12 | 12 |
| Total motility after thawing (%) | 14 | 44 | 29 | 48 | 12 | 57 |
| Sperm concentration after Swim-Up (M spz/mL) | 9 | 3 | 4 | 2 | 5 | 5 |
| Total motility after Swim-Up (%) | 64 | 64 | 54 | 65 | 53 | 81 |
| Sperm concentration before co-incubation (M spz/mL) | 4 | 2 | 2 | 3 | 4 | 4 |
| Total motility before co-incubation (%) | 60 | 52 | 63 | 75 | 58 | 62 |

Abbreviations: Cryo, Cryopreservation; Vitri, Vitrification
