## Supplementary Table S5 for "JUNO-Coated Beads as a Functional Assay to Capture and Characterize Fertilization-Competent Human Sperm"

**Supplementary Table S5. Donor information, biobank and laboratory thawed semen parameters, and sperm-bead binding assay outcomes.**

| Donor information | | | | Biobank thawing parameters[a] | | Laboratory thawing parameters[b] | | | | | | Experimental procedure | | | |
|---|---|---|---|---|---|---|---|---|---|---|---|---|---|---|---|
| SS | DA | BG | PF | PM (%) | NM (%) | SC_AT (M /mL) | TM_AT (%) | SC_ASU (M /mL) | TM_ASU (%) | SC_BI (M /mL) | TM_BI (%) | Spz_BJ | Spz_BC | Per_BJ (%) | Per_BC (%) |
| D048_310123_P11 | 21 | A+ | Yes | 80 | 8 | 11 | 46 | 6 | 69 | 6 | 65 | 3.61 | 0.23 | 90.9 | 16.1 |
| D003_240323_P02 | 27 | AB+ | Yes | 65 | 7 | 6 | 41 | 4 | 55 | 2 | 62 | 6.00 | 0.43 | 90.6 | 33.3 |
| D053_031022_P04 | 26 | O+ | Yes | 63 | 7 | 11 | 39 | 3 | 75 | 2 | 57 | 3.87 | 0.37 | 86.7 | 19.4 |
| D048_070223_P03 | 21 | A+ | Yes | 70 | 8 | 3 | 27 | 1 | 68 | 1 | 75 | 1.95 | 0.30 | 78.4 | 24.2 |
| D003_240323_P03 | 27 | AB+ | Yes | 65 | 7 | 7 | 50 | 1 | 56 | 3 | 75 | 12.37 | 0.26 | 96.7 | 19.4 |
| D053_031022_P02 | 26 | O+ | Yes | 63 | 7 | 7 | 43 | 2 | 88 | 2 | 78 | 7.23 | 0.45 | 87.9 | 24.3 |
| D005_011222_P02 | 28 | A+ | Yes | 65 | 12 | 5 | 33 | 1 | 60 | 1 | 72 | 3.97 | 0.19 | 82.9 | 19.4 |
| D005_011222_P04+P01 | 28 | A+ | Yes | 65 | 12 | 4.5 | 30 | 6 | 55 | 6 | 59 | 7.19 | 0.50 | 100.0 | 40.0 |
| D063_041023_P05+P04 | 22 | O+ | Yes | 67 | 8 | 4.5 | 28 | 2 | 50 | 5 | 44 | 5.55 | 0.51 | 100.0 | 35.0 |
| D048_190923_P05+P06 | 22 | A+ | Yes | 67 | 8 | 5 | 22 | 2 | 64 | 4 | 75 | 9.48 | 1.10 | 100.0 | 56.7 |
| D048_090124_P01 | 22 | A+ | Yes | 77 | 8 | 6 | 34 | 4 | 46 | 2 | 70 | 2.69 | 0.18 | 86.7 | 17.7 |
| D045_090124_P01 | 42 | O- | Yes | 78 | 9 | 7 | 44 | 3 | 64 | 2 | 52 | 17.58 | 0.90 | 100.0 | 60.0 |
| D089_040124_P01 | 19 | A+ | Yes | 86 | 6 | 5 | 48 | 2 | 65 | 3 | 75 | 11.66 | 0.48 | 100.0 | 41.9 |
| D048_090124_P02+P03 | 22 | A+ | Yes | 77 | 8 | 12 | 57 | 5 | 81 | 4 | 62 | 4.00 | 1.11 | 93.6 | 63.0 |
| D045_090124_P02 | 42 | O- | Yes | 78 | 9 | 5 | 38 | 6 | 61 | 6 | 60 | 3.91 | 0.31 | 95.2 | 52.7 |

| | | | | | | | | | | | | | | | |
|---|---|---|---|---|---|---|---|---|---|---|---|---|---|---|---|
| D045_090124_P03 | 42 | O- | Yes | 78 | 9 | 3 | 43 | 2 | 47 | 4 | 44 | 3.18 | 0.24 | 90.9 | 21.0 |
| D059_220923_P05 | 24 | O+ | Yes | 78 | 9 | 4 | 28 | 3 | 60 | 4 | 60 | 14.10 | 0.31 | 96.6 | 22.9 |
| D040_120923_P05 | 23 | A+ | Yes | 65 | 6 | 5 | 23 | 8 | 85 | 6 | 70 | 13.70 | 0.09 | 100.0 | 6.2 |
| D048_070223_P01 | 21 | A+ | Yes | 70 | 8 | 7 | 45 | 3 | 50 | 2 | 55 | 2.47 | 0.10 | 90.6 | 10.0 |
| D089_040124_P02 | 19 | A+ | Yes | 86 | 6 | 3 | 55 | 4 | 50 | 4 | 56 | 25.39 | 0.31 | 100.0 | 21.9 |
| D089_040124_P03 | 19 | A+ | Yes | 86 | 6 | 5 | 47 | 4 | 60 | 4 | 61 | 5.59 | 0.33 | 96.6 | 24.2 |
| D059_220923_P04 | 24 | O+ | Yes | 78 | 9 | 2 | 31 | 2 | 40 | 2 | 42 | 11.64 | 1.09 | 100.0 | 56.3 |
| D048_190923_P07 | 22 | A+ | Yes | 67 | 8 | 6 | 28 | 2 | 60 | 2 | 58 | 7.79 | 0.13 | 100.0 | 12.5 |
| D040_120923_P03 | 23 | A+ | Yes | 65 | 6 | 6 | 37 | 4 | 66 | 5 | 60 | 11.04 | 0.21 | 100.0 | 20.7 |
| D105_090224_P03 | 21 | A+ | Yes | 55 | 6 | 10 | 55 | 7 | 74 | 5 | 61 | 2.63 | 0.00 | 80.0 | 0.0 |
| D045_120424_P01 | 42 | O- | Yes | 70 | 9 | 8 | 33 | 4 | 61 | 5 | 60 | 7.73 | 0.24 | 100.0 | 20.6 |
| D092_020224_P01 | 27 | O+ | Yes | 73 | 7 | 6 | 35 | 4 | 60 | 4 | 58 | 3.39 | 0.12 | 91.2 | 12.1 |
| D089_211223_P01 | 19 | A+ | Yes | 72 | 6 | 6 | 65 | 3 | 70 | 4 | 69 | 5.63 | 0.42 | 100.0 | 39.4 |
| D003_141122_P01 | 27 | AB+ | Yes | 75 | 7 | 9 | 45 | 4 | 68 | 5 | 70 | 16.00 | 0.52 | 96.0 | 28.6 |
| D045_150224_P13+P05 | 42 | O- | Yes | 75 | 9 | 10 | 50 | 4 | 58 | 4 | 48 | 10.67 | 1.29 | 100.0 | 62.5 |
| D040_250923_P08+P09 | 23 | A+ | Yes | 75 | 6 | 4 | 44 | 3 | 76 | 4 | 68 | 2.63 | 1.13 | 96.7 | 58.3 |
| D003_141122_P02+P03 | 27 | AB+ | Yes | 75 | 7 | 10 | 60 | 3 | 64 | 4 | 79 | 7.00 | 1.10 | 92.3 | 33.3 |
| D096_220224_P03+P04 | 30 | A+ | Yes | 67 | 7 | 7 | 32 | 2 | 51 | 4 | 50 | 6.13 | 0.26 | 90.0 | 22.6 |
| D045_150224_P06 | 42 | O- | Yes | 75 | 9 | 6 | 48 | 3.6 | 68 | 4 | 66 | 16.64 | 1.87 | 100.0 | 58.1 |
| D089_010923_P07 | 19 | A+ | Yes | 80 | 6 | 27 | 43 | - | - | 13 | 52 | 8.73 | 0.08 | 100.0 | 8.3 |
| D108_270224_P01+P02 | 34 | A+ | Yes | 55 | 6 | 11 | 50 | - | - | 4 | 41 | 4.80 | 0.21 | 94.0 | 21.0 |
| D040_031023_P05 | 27 | AB+ | Yes | 72 | 6 | 10 | 50 | - | - | 13 | 63 | 5.06 | 1.29 | 100.0 | 61.3 |
| D003_080923_P02 | 23 | A+ | Yes | 67 | 7 | 5 | 59 | - | - | 5 | 46 | 19.60 | 1.42 | 100.0 | 69.2 |
| D075_70224_P05 | 31 | A+ | Yes | 72 | 7 | 3 | 23 | - | - | 8 | 26 | 3.03 | 2.32 | 88.6 | 24.4 |
| D015_281024_P07 | 34 | O+ | No | 80 | 9 | 4 | 26 | - | - | 4 | 37 | 8.90 | 1.41 | 96.6 | 40.6 |
| D003_080923_P03 | 29 | AB+ | Yes | 67 | 7 | 3 | 53 | - | - | 3 | 56 | 5.19 | 0.90 | 96.8 | 44.8 |

| | | | | | | | | | | | | | | | |
|---|---|---|---|---|---|---|---|---|---|---|---|---|---|---|---|
| D133_131124_P02 | 29 | O+ | No | 65 | 9 | 3 | 55 | - | - | 4 | 78 | 8.67 | 0.88 | 100.0 | 38.5 |
| D108_070524_P07 | 35 | A+ | Yes | 62 | 6 | 3 | 66 | - | - | 4 | 46 | 18.25 | 0.39 | 100.0 | 32.3 |
| D032_140224_P01 | 26 | A+ | Yes | 65 | 9 | 6 | 35 | - | - | 12 | 41 | 15.93 | 1.70 | 100.0 | 63.3 |
| D015_281024_P02 | 34 | O+ | No | 80 | 9 | 12 | 50 | - | - | 13 | 44 | 4.20 | 1.41 | 93.3 | 55.6 |
| D108_070524_P08 | 35 | A+ | Yes | 62 | 6 | 12 | 54 | - | - | 12 | 44 | 6.50 | 1.00 | 100.0 | 45.5 |
| D108_070524_P09 | 35 | A+ | Yes | 62 | 6 | 7 | 75 | - | - | 5 | 38 | 11.07 | 0.44 | 96.4 | 32.4 |
| D003_080923_P05 | 29 | AB+ | Yes | 67 | 7 | 8 | 57 | - | - | 5 | 45 | 7.87 | 0.44 | 100.0 | 25.0 |
| D133_131124_P03 | 29 | O+ | No | 65 | 9 | 19 | 49 | - | - | 18 | 33 | 7.67 | 1.79 | 90.0 | 48.5 |
| D032_140224_P03 | 26 | A+ | Yes | 65 | 9 | 12 | 57 | - | - | 9 | 29 | 9.55 | 0.09 | 100.0 | 8.8 |
| D015_281024_P08 | 34 | O+ | No | 80 | 9 | 16 | 48 | - | - | 8 | 30 | 3.23 | 0.19 | 90.3 | 18.8 |

SS, Semen sample (Donor ID_ejaculate_straw); DA, Donor Age (years); BG, Blood Group (ABO and Rh); PF, Proven fertility; PM, Progressive motility (%); NM, Normal morphology (%); SC_AT, Sperm concentration after thawing (M spz/mL); TM_AT, Total motility after thawing (%); SC_ASU, Sperm concentration after Swim-Up (M spz/mL); TM_ASU, Total motility after Swim-Up (%); SC_BI, Sperm concentration before co-incubation (M spz/mL); TM_BI, Total motility before co-incubation (%); Spz_BJ, Mean number of spz bound to BJUNO; Spz_BC, Mean number of spz bound to Bcontrol; Per_BJ, BJUNO with ≥ 1 spz bound (%); Per_BC, Bcontrol with ≥ 1 spz bound (%); -, no data obtained.

[a]Semen parameters provided by the biobank company, assessed using a thawing test straw.
[b]Laboratory-assessed semen parameters after thawing the experimental straws.
