## Supplementary Figure S1 for "JUNO-Coated Beads as a Functional Assay to Capture and Characterize Fertilization-Competent Human Sperm"

**A**

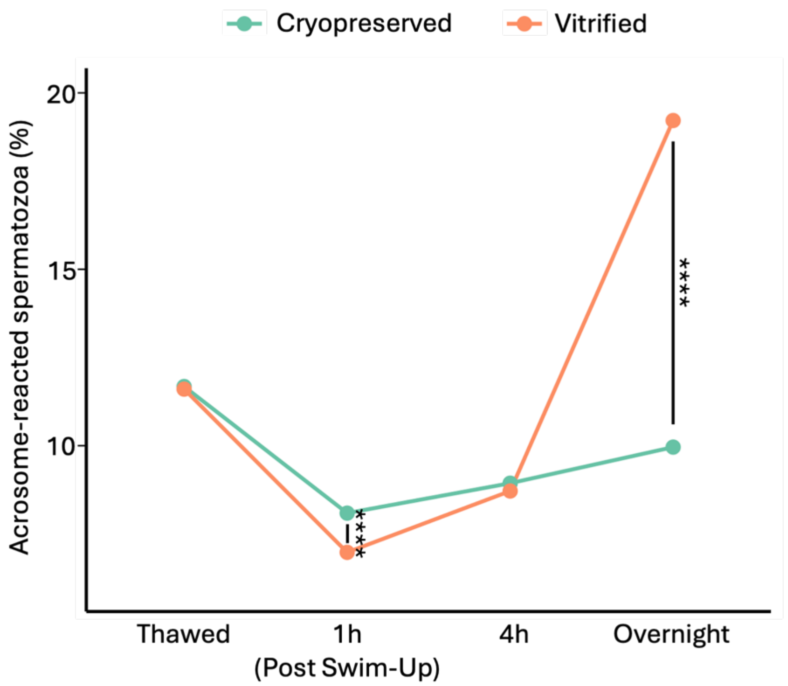


**B**

| **Incubation time** | **Cryopreserved** (%) | **Vitrified** (%) | ***p* value** |
|---|---|---|---|
| Thawed | 11.7 | 11.6 | 0.7837 |
| 1h (Post Swim-Up) | 8.1 | 7.0 | <0.0001 |
| 4h | 8.9 | 8.7 | 0.5271 |
| Overnight | 10.0 | 19.2 | <0.0001 |

**Supplementary Fig S1**. **Freezing method influences sperm acrosome reaction capacity**. **A**. Percentage of acrosome-reacted sperm cells over time under capacitating conditions at different time point, including immediately after thawing, sperm selection for 1 hour (Swim-Up), 4 hours and overnight incubation. ****$p \leq 0.0001$. **B**. Numerical values corresponding to panel A. Flow cytometry analysis reveals that vitrification better preserves the acrosome reaction capacity after overnight incubation.
