## Supplementary Figure S2 for "JUNO-Coated Beads as a Functional Assay to Capture and Characterize Fertilization-Competent Human Sperm"

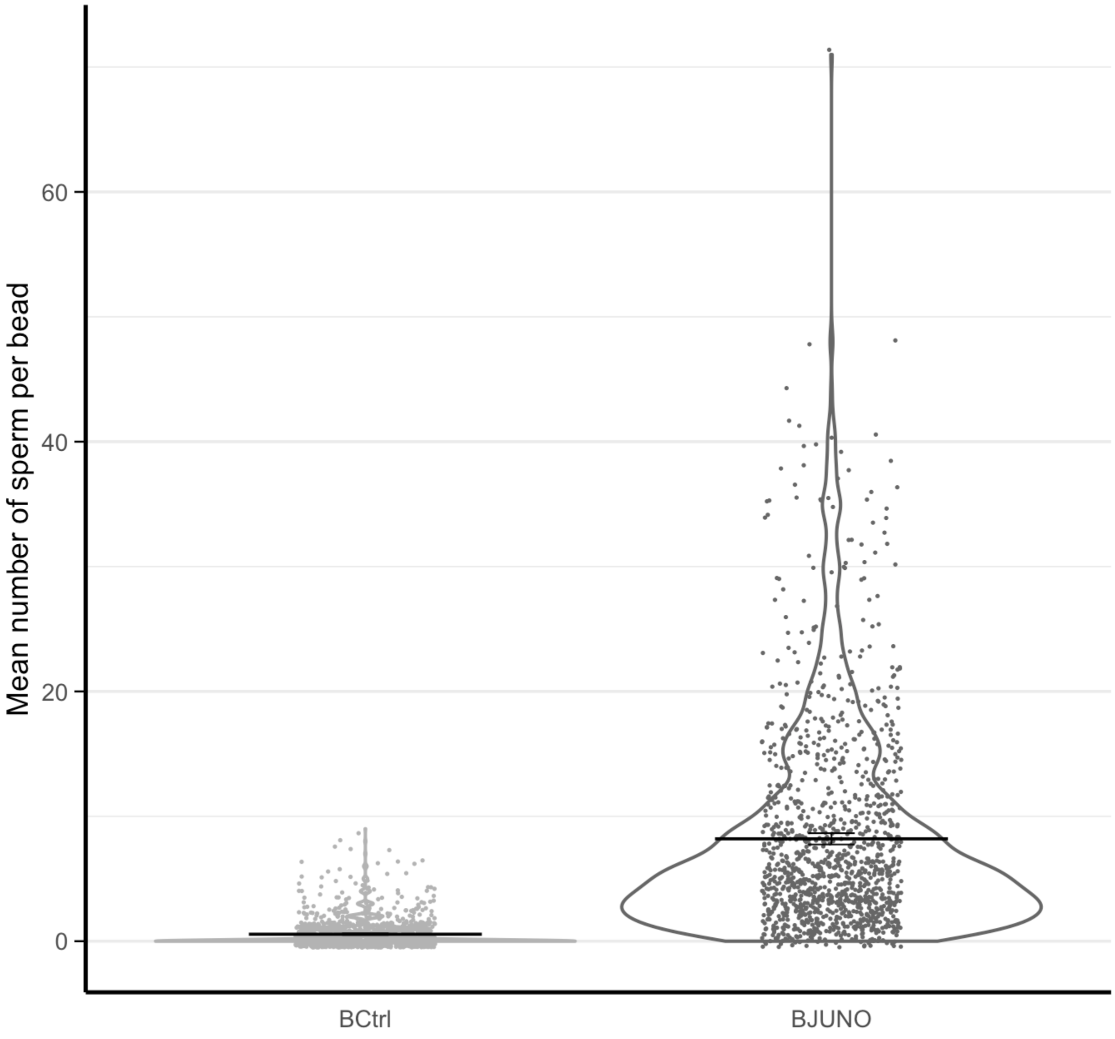


**Supplementary Fig S2**. **Human spermatozoa bind specifically to JUNO-coated beads**. Numerical values corresponding to panel A. Flow cytometry analysis reveals that vitrification better preserves the acrosome reaction capacity after overnight incubation.
